# Reflective imaging improves resolution, speed, and collection efficiency in light sheet microscopy

**DOI:** 10.1101/154807

**Authors:** Yicong Wu, Abhishek Kumar, Corey Smith, Evan Ardiel, Panagiotis Chandris, Ryan Christensen, Ivan Rey-Suarez, Min Guo, Harshad D. Vishwasrao, Jiji Chen, Jianyong Tang, Arpita Upadhyaya, Patrick J. La Riviere, Hari Shroff

**Affiliations:** Section on High Resolution Optical Imaging, National Institute of Biomedical Imaging and Bioengineering, National Institutes of Health, Bethesda, MD, USA; Dept. of Radiology, University of Chicago, Chicago, IL, USA; Biophysics Program, University of Maryland, College Park, MD, USA; Advanced Imaging and Microscopy Resource, National Institutes of Health, Bethesda, MD, USA; JT Scientific Consulting LLC, North Potomac, MD, USA; Department of Physics and Institute of Physical Science and Technology, University of Maryland, College Park, MD, USA; Whitman Center, Marine Biological Laboratory, Woods Hole, MA, USA

## Abstract

Light-sheet fluorescence microscopy (LSFM) enables high-speed, high-resolution, gentle imaging of live biological specimens over extended periods. Here we describe a technique that improves the spatiotemporal resolution and collection efficiency of LSFM without modifying the underlying microscope. By imaging samples on reflective coverslips, we enable simultaneous collection of multiple views, obtaining 4 complementary views in 250 ms, half the period it would otherwise take to collect only two views in symmetric dual-view selective plane illumination microscopy (diSPIM). We also report a modified deconvolution algorithm that removes the associated epifluorescence contamination and fuses all views for resolution recovery. Furthermore, we enhance spatial resolution (to < 300 nm in all three dimensions) by applying our method to a new asymmetric diSPIM, permitting simultaneous acquisition of two high-resolution views otherwise difficult to obtain due to steric constraints at high numerical aperture (NA). We demonstrate the broad applicability of our method in a variety of samples of moderate (< 50 μm) thickness, studying mitochondrial, membrane, Golgi, and microtubule dynamics in single cells and calcium activity in nematode embryos.

## Introduction

Modern light-sheet fluorescence microscopy (LSFM) combines high speed, optical sectioning, diffraction-limited spatial resolution and low phototoxicity. The consolidation of these attributes makes LSFM an attractive choice for volumetric, time-lapse (4D) imaging in living samples spanning spatial scales from the subcellular^1^ to organismal^2^, and temporal scales from milliseconds^3^ to days^4^. Continued technical development has proved essential in applying LSFM to biology^5^.

Many implementations of LSFM serially acquire multiple specimen views and computationally fuse them, compensating for anisotropic spatial resolution^6^ or specimen-induced scatter^7,8^ that plague any individual view. While powerful, serial multiview imaging entails significant additional cost in temporal resolution and phototoxicity, as acquisition time and illumination dose increase with every additional view. At low NA, temporal resolution in serial multiview LSFM can be dramatically improved by using multiple objectives to simultaneously image perpendicular views with phase-shifted confocal line detection^2^. To remove crosstalk between views, the phase offset was set at a relatively large value (tens of μm), thus limiting the method to large samples. Furthermore, imaging at higher NA (suitable for smaller samples such as single cells or nematode embryos) is confounded by the relatively short working distance of the detection lenses, since apposing multiple bulky lenses and a sample is difficult or impossible due to steric constraints.

We demonstrate a simple and effective solution to these problems by using commercially available reflective coverslips to provide additional specimen views, and a modified joint deconvolution algorithm to remove epifluorescence contamination and fuse the views. In a symmetric dual-view LSFM setup (diSPIM, 0.8 NA/0.8 NA detection), reflective coverslips allow four complementary views to be acquired simultaneously, doubling light collection and speed relative to conventional dual-view imaging. These enhancements enable the study of rapid calcium dynamics in nematode embryos and the capture of fast membrane and mitochondrial dynamics in live cells at isotropic spatial resolution (^~^330 nm in all 3 spatial dimensions), uncovering transient events otherwise blurred by diffraction or motion. In a new asymmetric diSPIM, our technique further improves spatial resolution (to ^~^260 nm laterally and ^~^300 nm axially) and collection efficiency (to 44% of the available solid angle compared to only ^~^20% with conventional symmetric diSPIM), as reflection allows us to obtain complementary 1.1 NA views that are otherwise difficult to collect due to steric constraints. We illustrate the power of our approach by imaging dynamics of microtubules, membranes, Golgi, and mitochondria in single cells and calcium flux in nematode embryos. In all cases, reflective imaging boosts information content of images, improving the spatiotemporal resolution of the underlying microscope technology.

## Results

### Using reflection for multiview LSFM imaging

Reflection has previously been used to introduce LSFM excitation to the sample, cleverly permitting the use of high-NA detection objectives in single-view imaging^9-11^. Although reflection has also been used in epifluorescence imaging to collect additional specimen views ^12,13^ (albeit without view fusion), we are unaware of any similar efforts in LSFM. We reasoned that if samples were placed on reflective substrates, and light-sheet illumination and detection optics introduced appropriately (e.g. with the diSPIM geometry), two significant advantages would accrue. First, the light sheet would reflect off the coverslip, creating a second orthogonal light sheet, allowing for simultaneous rather than sequential diSPIM imaging. Second, the fluorescence from each light sheet that would otherwise be lost through the coverslip would reflect off the coverslip and be imaged by the cameras. Effectively, the fluorescence reflection would duplicate the imaging geometry, thereby creating a virtual image of the sample, illumination, and detection pathways on the underside of the coverslip (Fig. 1a). The advantage of this reflection is that it enables each objective in our diSPIM implementation of LSFM to capture 2 orthogonal views simultaneously (A and mirror image B’; B and mirror image A’, Fig. 1a) as the sample is translated through direct and reflected light sheets. We confirmed this intuition by imaging live EGFP-histone labeled Caenorhabditis elegans embryos deposited on commercially available reflective coverslips in a stage-scanning diSPIM with synchronized rolling shutter confocal slit detection^14^ (Fig. 1b, c, **Supplementary Video 1**). In principle, these 4 views can be registered and fused to improve: (i) spatial resolution (especially axially, since the poor axial resolution in any single view can be compensated by the much better lateral resolution of its perpendicular partner^6^), (ii) signal-to-noise ratio (since fluorescence that is usually wasted can now be collected via the reflection, enabling collection of 4 views instead of the usual 2), and (iii) imaging speed (since both cameras can record LSFM data simultaneously, instead of the usual alternating duty cycle in conventional diSPIM)^6^.

**Fig. 1.**
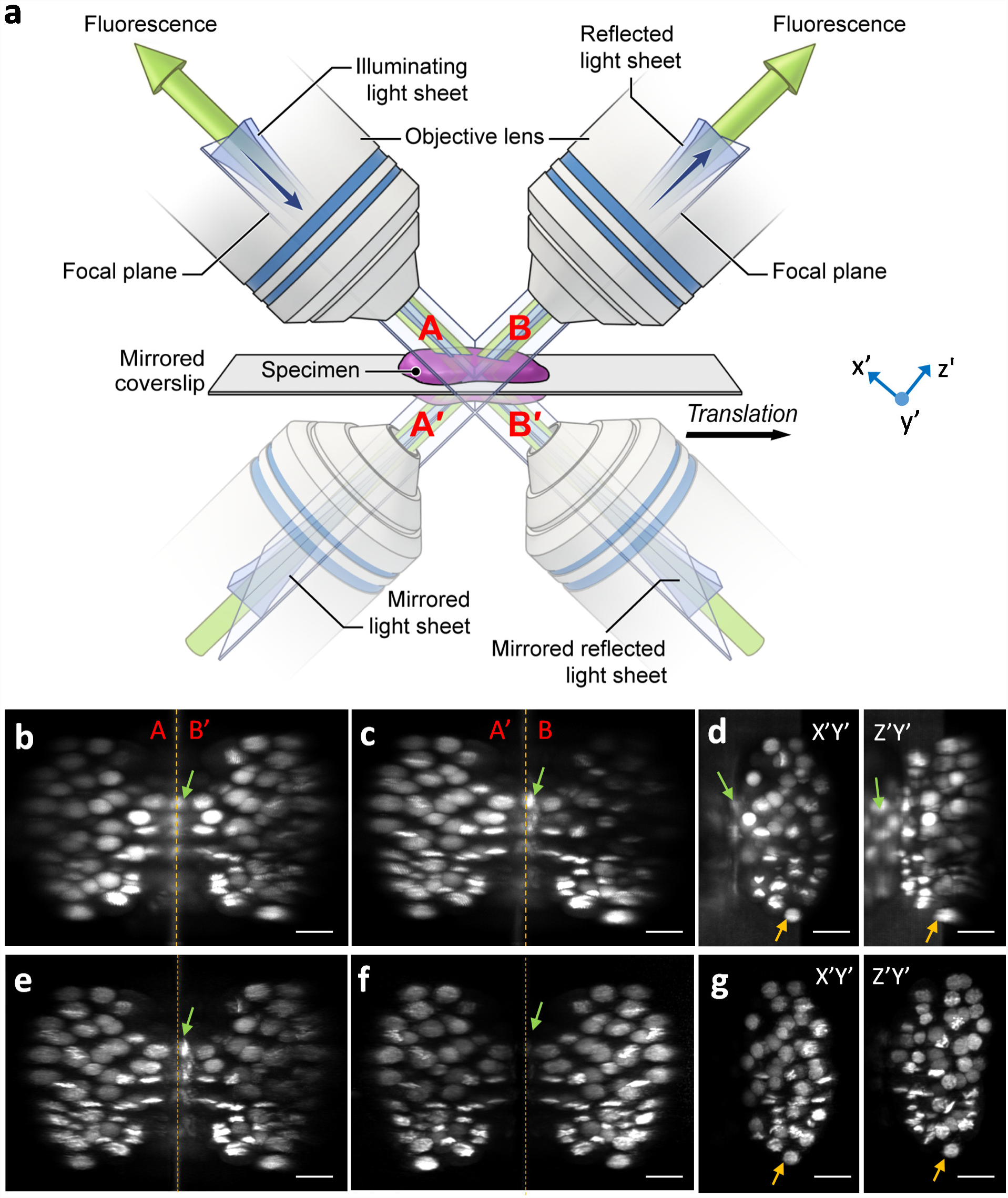
Reflective imaging in light sheet microscopy. **a)** Schematic indicating reflective light sheet microscopy (symmetric diSPIM) geometry. Sample, objectives, and light sheets are indicated above mirrored coverslip, ‘virtual’ copies shown below the coverslip. In this example, the light sheet is introduced from the left, and reflected by the mirror. Four views, indicated A, A’ (mirror image of A), B (produced by reflected light sheet) and B’ (mirror image of B) are simultaneously obtained as the sample is translated through the stationary light sheets. In other words, A’ and B’ appear to originate from virtual sources below the mirrored coverslip. The reference coordinate system (from the perspective of the right hand objective) is also indicated. Maximum intensity projections of raw volumetric views of EGFP-histone labeled nuclei in a live nematode embryo, as collected from either objective as the sample is translated through the stationary light sheets are shown in **b, c**. Data from view A is transformed (deskewed) so that it is shown as if the light sheets are scanned through a stationary sample, **d**. Naïve deconvolution results in the reconstruction shown in **e**, but better modeling of the imaging process results in the reconstructions shown in **f** (shown in **g** from the same perspective as in **d**). Note that epifluorescence contamination (green arrows) is markedly reduced in **f, g** (compare to **b-e**) and resolution isotropy restored (compare **d, g**, especially nuclei highlighted with orange arrows). Scale bars: 10 μm. See also **Supplementary Videos 1-2.**

However, each view is contaminated by substantial background caused by out-of-focus fluorescence excited by the light sheets, since illumination is no longer entirely coincident with each detection plane. This contamination is most severe near the coverslip where the two light sheets cross, and is evident when viewing the raw data (Fig. 1b, c) and when viewing the data from the perspective of the objective (i.e., deskewed, Fig. 1d). Registering all 4 views and naively fusing them using joint-deconvolution with detection point spread functions appropriate for each lens failed to eliminate the background contamination (Fig. 1e, **Supplementary Video 2**), not surprisingly, because such an approach fails to account for spatially-varying out-of-focus fluorescence.

We hypothesized that carefully accounting for the background in our deconvolution model would improve the fusion. To this end we (i) modeled the imaging geometry, accounting for all 4 LSFM views and associated spatially varying epifluorescence background by considering the sample, the illumination, and their reflections; (ii) modeled the spatially varying imaging process, including rolling-shutter slit detection; (iii) modified Richardson-Lucy deconvolution for such spatially varying imaging, computing appropriate forward and transpose operators that relate object and image formation; and (iv) registered the raw image volumes, adapted our modified Richardson-Lucy deconvolution algorithm for use with a parallelized, graphics processing unit (GPU) implementation to speed processing, and applied the algorithm to reconstruct the data. These steps are described in detail in **Online Methods** and **Supplementary Notes 1-3**. Applying them to the raw data greatly reduced epifluorescence contamination (Fig. 1f) and restored resolution isotropy (Fig. 1g, **Supplementary Video 2**). Also, selecting the “best” data from each view could be used to reduce the effect of scattering that degrades any individual view (**Supplementary Fig. 1, Methods**).

Reflective imaging and subsequent deconvolution enabled observation of high-speed phenomena that would be otherwise masked by diffraction^15^ or motion blur^6^ in other LSFM implementations. For example, we were able to “catch” calcium waves at isotropic spatial resolution when imaging a freely moving nematode embryo expressing GCaMP3 under the control of the ***myo-3*** promoter (Fig. 2a, **Supplementary Video 3**). We attained a continuous volumetric imaging rate of 2.86 Hz in the reflective diSPIM, in part by forgoing synchronized rolling-shutter confocal slit detection (**Methods**). Although using a rolling confocal shutter does reduce background (**Supplementary Video 4**), our modified deconvolution algorithm enabled imaging with acceptable signal-to-background even without the shutter (**Supplementary Fig. 2, Methods**). Using the reflective diSPIM without the rolling shutter, we observed similar calcium dynamics as those obtained with earlier single view LSFM^16^, but with much better axial resolution (Fig. 2b, **Supplementary Fig. 3**). The combination of high speed and resolution isotropy enabled us to easily define events in the axial view (Fig. 2b), such as the onset and subsequent spread of a bidirectional wave, occurring within one second (Fig. 2c). In another example, we observed the rise and decay of dF/F (change in fluorescence intensity over baseline) within three closely spaced muscle bundle positions (Fig. 2d, e). The 2-fold higher speed of reflective diSPIM relative to conventional diSPIM (i.e. using glass coverslips) allowed us to separate the dynamics temporally, revealing that the event in one area lagged the others by ~350 ms (Fig. 2e). Such changes would have been blurred in conventional diSPIM, obscuring underlying differences in onset times. Furthermore, the 4 raw views simultaneously collected during each “pass” of the sample through the direct and reflected light sheets proved useful in confirming the presence of true calcium transients, since these events were visible in each view (**Supplementary Fig. 3**).

**Fig. 2.**
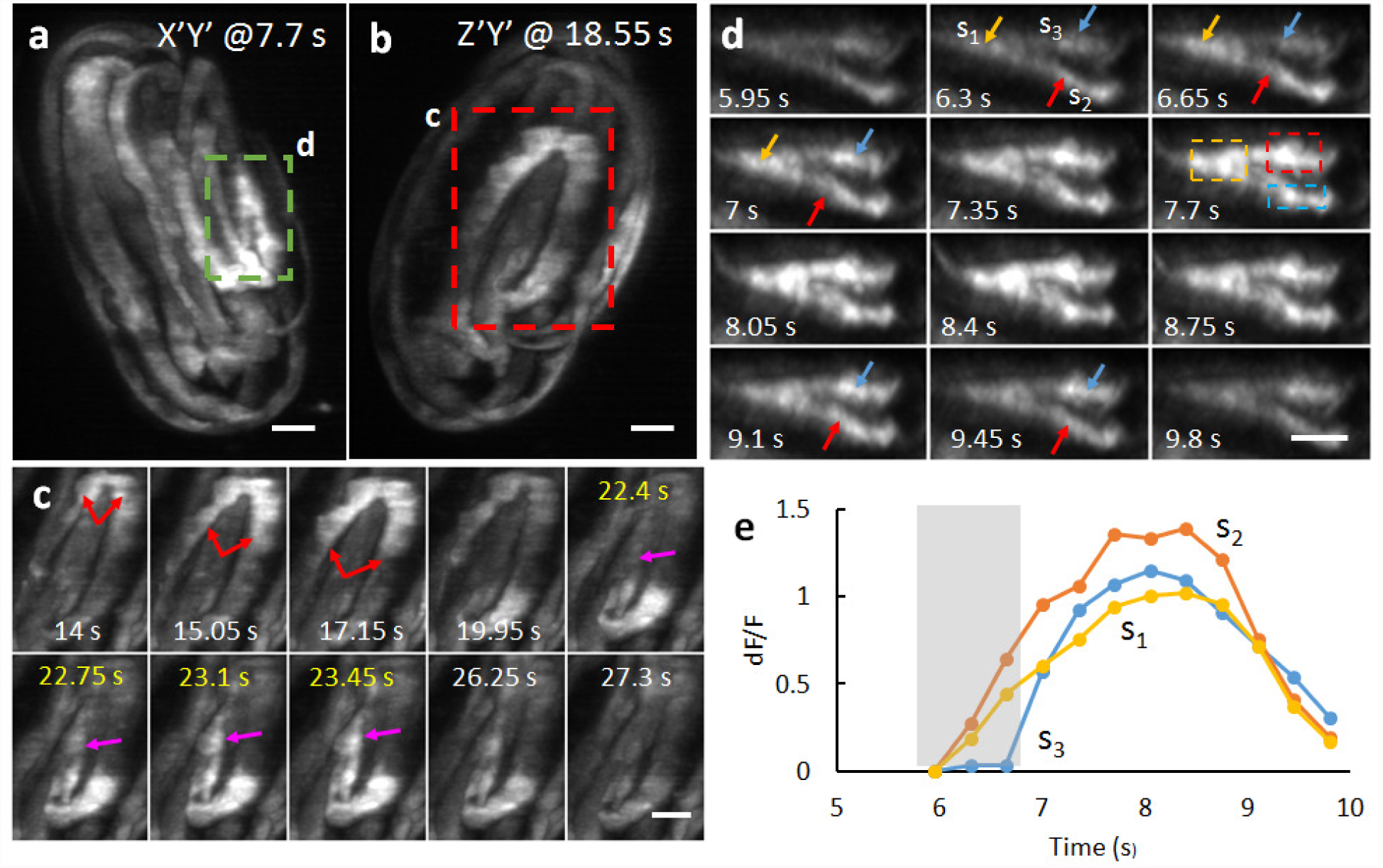
Single-pass isotropic imaging with reflective diSPIM reveals rapid muscular calcium flux within 3-fold nematode embryo. **a, b)** Maximum intensity projections of 3-fold embryo expressing GCaMP3 from a *myo-3* promoter at two different time points and from lateral (**a**) and axial (**b**) views. **c**) Higher magnification view of red dashed rectangular region in **b)** at indicated time points. Red arrows illustrate a spreading wave that appears to propagate bidirectionally. Purple arrow indicates a spreading event that appears to form and propagate within a second; yellow headings indicate successive timepoints (volumes acquired every 350 ms). **d)** Higher magnification view of green dashed rectangular region in **a)**. Twelve successive time points are shown, indicating onset and eventual disappearance of intensity transient. Three areas (s1, s2, s3) are highlighted with yellow, red, and blue arrows. Intensities from each area (quantified from dashed rectangular regions) are shown in **e**). Grey bounding area highlights differential rise in intensity in areas s1, s2 vs. s3, which would be masked if the data were acquired more slowly. See also **Supplementary Videos 3-4, Supplementary Fig. 3**. All scale bars: 5 μm.

To demonstrate the usefulness of reflective diSPIM imaging in 2-color applications, where subcellular resolution is required, we imaged U2OS cells transfected with TurboGFP-Lck (labeling the plasma membrane, endocytic machinery, and Golgi compartment) and Tom20-mApple (labeling the outer mitochondrial membrane), collecting 300 2-color volumes (Fig. 3a, **Supplementary Video 5**). Our high temporal resolution (one 2-color volume every 5 s) facilitated inspections of protrusive plasma membrane (Fig. 3b) and mitochondrial dynamics (e.g., fusion and fission, Fig. 3c) both near and far (Fig. 3d-f) from the coverslip surface. As in conventional diSPIM, resolution isotropy was maintained throughout the cell volume, as revealed by examination of membrane-enclosed vesicles (Fig. 3g). Comparisons of reconstructions to the raw data confirmed the validity of our imaging model and underscored the importance of epifluorescence removal and joint deconvolution in restoring fine details in the data (compare Fig. 3a to Fig. 3h, Fig. 3c to Fig. 3i and Fig. 3g to Fig. 3j, **Supplementary Video 6**).

**Fig. 3.**
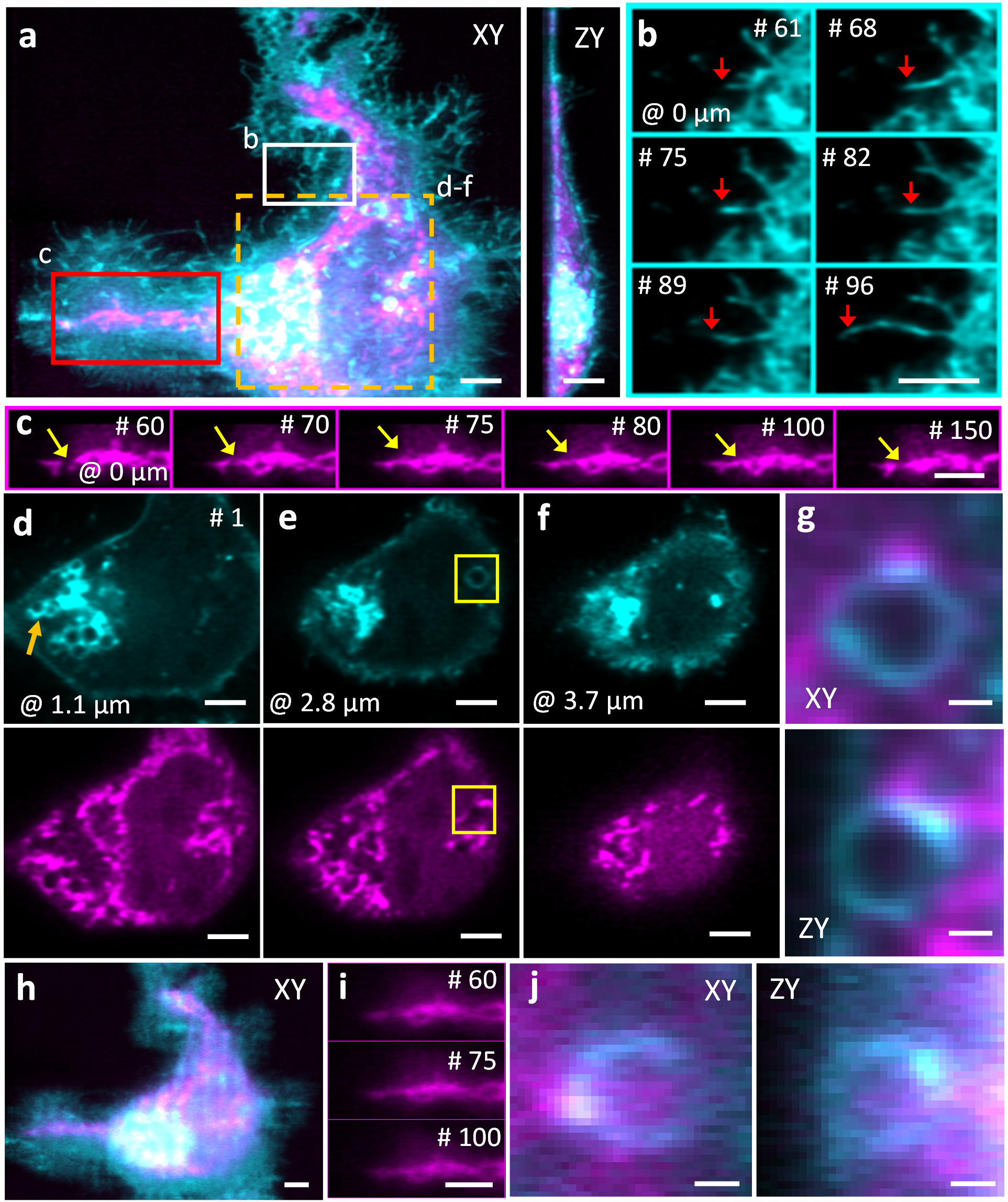
Reflective diSPIM imaging reveals mitochondrial and membrane dynamics over hundreds of imaging volumes. **a)** Maximum intensity projection of U2OS cell labeled with TurboGFP-Lck (cyan, staining the plasma membrane, endocytic machinery, and Golgi compartment) and Tom20-mApple (magenta, staining the outer mitochondrial membrane). Lateral (left) and axial (right) projections are shown, coordinates are defined with respect to the coverslip. **b)** Higher magnification lateral view of white rectangular region in **a)**, emphasizing protrusive dynamics at the plasma membrane, most likely representing filopodia. Overall growth of the filament is evident from volumes 61-96 (tip of filament highlighted with red arrow). **c)** Higher magnification views of the red rectangular region in **a)**, emphasizing mitochondrial dynamics. Mitochondria show fusion (volume 75) and fission (volume 150); yellow arrow marks the site of these events. **d**-**f)** Selected imaging planes at indicated axial location in the first volume, top row showing membranous structures and bottom row showing mitochondria. Membranes and mitochondria are concentrated at the left edge of the cell, near the Golgi system (yellow arrow in **d**). **g)** Higher magnification view of endocytic vesicle and surrounding region populated with mitochondria marked within yellow rectangle in **e**). Lateral (top) and axial (bottom) slices are shown to emphasize resolution isotropy. **h-j)** Same as **a, c, g** but raw data from view A. Data are selected from an imaging series spanning 300 volumes (imaging duration 25 min). See also **Supplementary Videos 5-6**. Scale bars: 5 μm (except **g, j,** 1 μm).

### Reflective diSPIM imaging at higher spatial resolution

Our success in applying reflecting imaging to the symmetric 0.8 NA / 0.8 NA diSPIM motivated us to apply the same concept to LSFM with higher NA detection. As a first step, we designed and constructed an asymmetric diSPIM with 1.1 NA and 0.71 NA objectives for illumination/collection (Fig. 4a, **Supplementary Fig. 4, Methods**), using fiber-coupled laser illumination in commercially available scanheads^17^ and two sCMOS cameras for detection. Since the overall NA of the asymmetric system is larger, collection efficiency increased ^~^1.5 fold (Fig. 4b), and volumetric resolution ^~^1.4-fold relative to the symmetric diSPIM after joint deconvolution (**Methods**, Fig. 4c, **Supplementary Table 1**). Although the different collection objectives in the asymmetric design produce an asymmetric PSF with slightly worse axial resolution than the symmetric case (0.71/0.8 = 0.89 x), areal resolution in the focal plane is improved (1.1/0.8)^2 = 1.89 x, an advantage that was obvious when visualizing immunolabeled microtubules in fixed U2OS cells (Fig. 4d). When imaging Alexa Fluor 488 nm phalloidin-labeled actin in U2OS cells plated on glass coverslips (Fig. 4f), the asymmetric system enabled us to visualize fine filaments throughout the 5.2 μm-thick volume (**Supplementary Video 7**), resolving filaments separated by ^~^260 nm (Fig. 4h, **Supplementary Fig. 5**). The overall resolution improvement was also evident when comparing images of live nematode embryos deposited on glass coverslips and imaged with either symmetric or asymmetric systems (**Supplementary Fig. 6**).

**Fig. 4.**
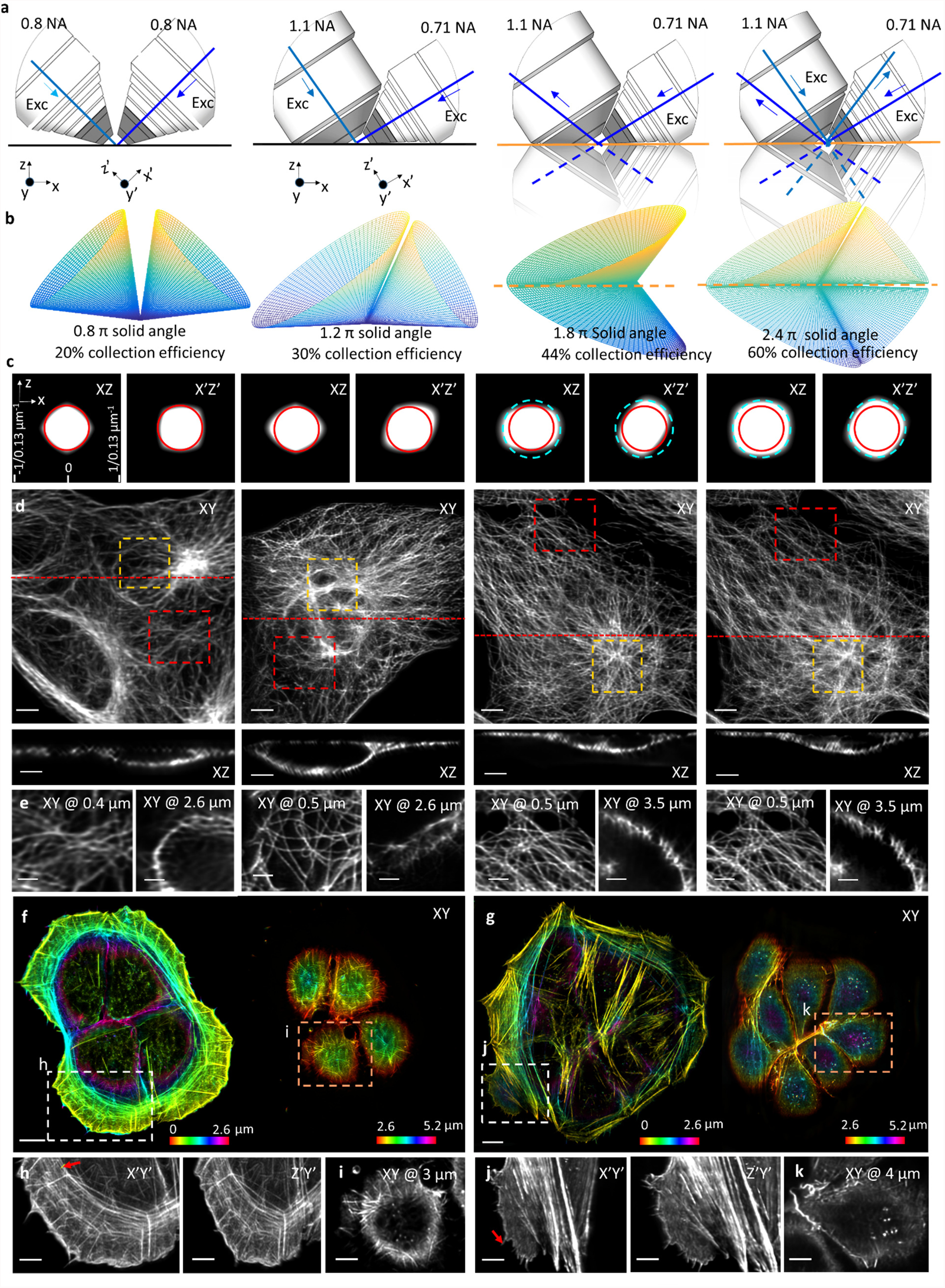
Reflective imaging at high NA boosts resolution and collection efficiency. **a)** Comparative schematics showing from left to right, 0.8 NA/0.8 NA dual-view on glass coverslip; 1.1 NA/0.7 NA dual-view on glass coverslip; 1.1 NA/1.1 NA dual-view on reflective coverslip; 1.1 NA/0.7 NA quadruple-view on reflective coverslip. Excitation light sheets are represented as blue lines, with arrows indicating the direction of propagation. Virtual light sheets are shown as dashed blue lines, virtual objectives are transparent. Primed (perspective from detection objective) and unprimed (from coverslip) coordinate axes are also shown. **b)** Collection cones for each detection lens geometry corresponding to **a)**, including total solid angle and corresponding collection efficiency. **c)** Cross sections through optical transfer functions (OTFs) in primed and unprimed coordinate systems, with diffraction limit indicated for 0.8 NA detection (red circle) and 1.1 NA detection (blue dashed circle). **d)** Lateral maximum intensity projections of images of Alexa Fluor 488 immunolabeled microtubules in fixed U2OS cells (top row) derived from each imaging system are shown, as are axial slices (bottom row) corresponding to red dotted lines. Higher magnification single plane views at indicated axial depths **(e);** closer (red dashed rectangular regions in **d)**) and farther (yellow rectangular regions in **d)**) from the coverslip are also shown. Examples of Alexa Fluor 488 phalloidin staining actin in fixed U2OS cells are also provided for 1.1 NA / 0.7 NA dual-view geometry on glass coverslip (**f**) and 1.1 / 1.1 NA dual-view geometry on reflective coverslip **(g)** are also shown, with axial depth from coverslip indicated with colorbar. Higher magnification views in primed **(h, j)** and unprimed **(i, k)** coordinate systems corresponding to dashed rectangular regions in **f, g** are also provided. See also **Supplementary Fig. 5** and **Supplementary Videos 7-8**. Scale bars: 5 μm **d**; 3 μm **e**; 10 μm **f**, **g**; 5 μm **h-k**.

Next, we investigated the performance of the system on reflective coverslips. When illuminating with the 0.71 NA lens and collecting via the 1.1 NA lens, two high-NA views oriented ^~^112 degrees apart were collected simultaneously when sweeping the sample through incident and reflected light sheets (Fig. 4a). This configuration not only doubled the imaging speed relative to conventional diSPIM imaging, but also enabled the collection of two high-NA views, which would be expensive and difficult to implement sterically with two physical objectives. Collection efficiency improved to ^~^44% and volumetric resolution to 260 × 260 × 300 nm (Fig. 4b, c). Relative to data obtained with the symmetric diSPIM on glass coverslips, we obtained sharper images of immunolabeled microtubules, after removing epifluorescence contamination and performing joint deconvolution (**Methods**), especially evident when viewed from the perspective of the coverslip (Fig. 4d, e, **Supplementary Fig. 5**). Images of phalloidin-labeled actin (Fig. 4g, j, k, **Supplementary Fig. 5, Supplementary Video 8**) were similar to those obtained with the asymmetric system on glass coverslips.

Since the direct and reflected light sheets are not orthogonal in the asymmetric diSPIM, collection of four light sheet views is not possible using illumination only from the 0.71 NA lens. However, alternating excitation and detection between the two objectives provides the requisite views **(Fig. 4a). Collection efficiency through both lenses in one such imaging cycle improved to ~60% (Fig. 4b)**, and resolution improved to near isotropic 260 × 260 × 280 nm, after removal of epifluorescence contamination and performing joint deconvolution (Fig. 4c). Images of microtubules (Fig. 4d-e, **Supplementary Fig. 5**) were marginally improved, but the extra illumination, acquisition time and (especially) computational cost associated with such improvement are considerable (**Supplementary Table 1**). For the remainder of the studies in this work, we thus used only the 2 high-NA views collected when illuminating with the 0.71 NA lens.

### Imaging subcellular dynamics and signal transduction at high spatiotemporal resolution with two high NA views

To demonstrate the potential of direct and reflected 1.1 NA views to visualize extended, 3D dynamics at high spatiotemporal resolution, we applied the technique to study a variety of biological processes in single cells and nematode embryos (Fig. 5, **Supplementary Videos 9-12**).

**Fig. 5.**
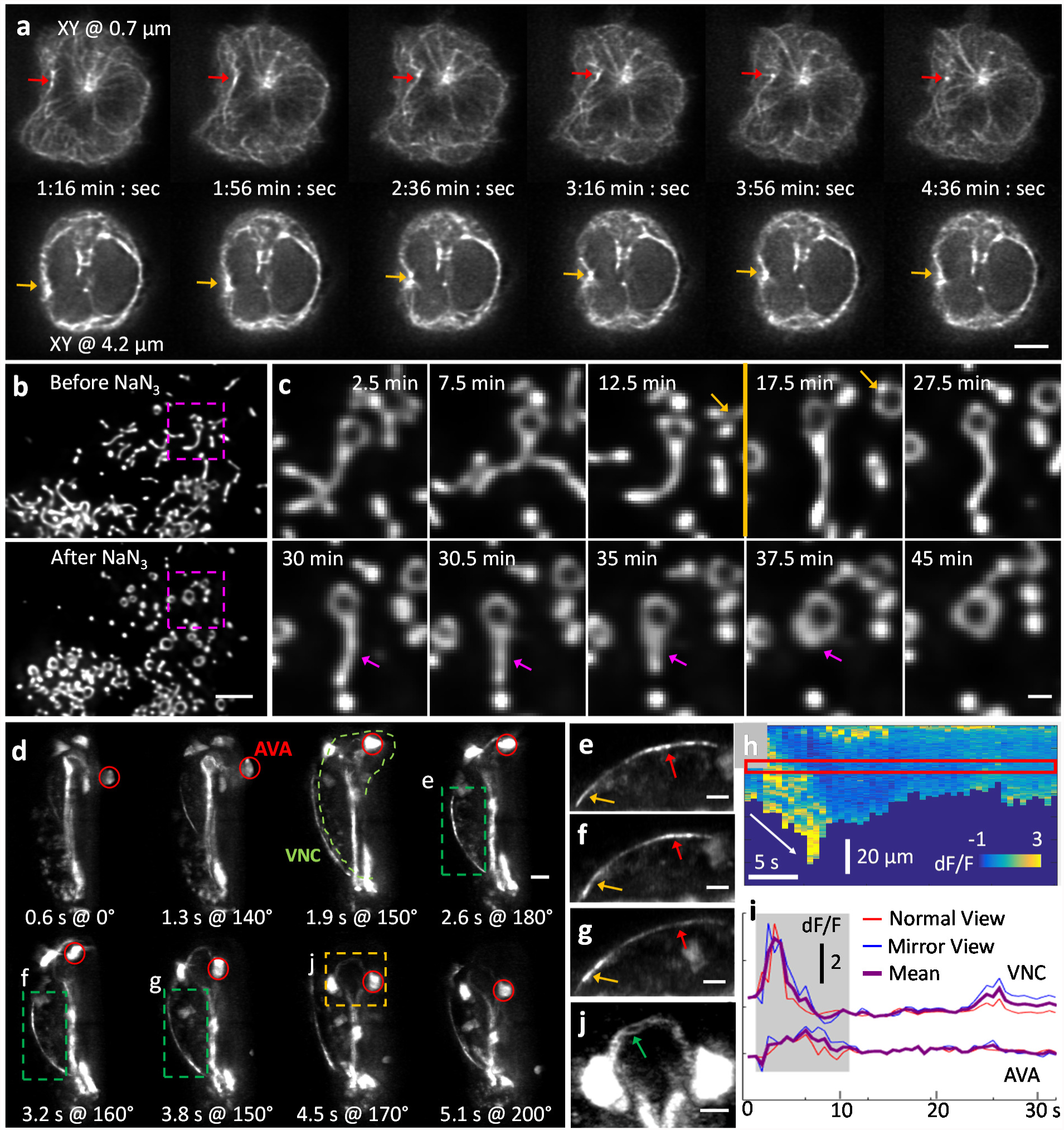
Volumetric time-lapse imaging with two high NA views enables investigation of subcellular dynamics at high spatiotemporal resolution. **a)** Single planes at indicated depth from coverslip, showing dynamics of microtubules in Jurkat T cells expressing 3XEGFP-EMTB. Red arrow in upper series highlights inward and subsequent outward displacement of microtubule bundles at base of cell, yellow arrow in lower series shows contemporaneous deformation of the left nuclear edge nucleus. **b)** Maximum intensity projections showing response of GFP- labeled mitochondria in U2OS cells to NaN3, an inhibitor of cytochrome c oxidase. **c)** Higher magnification views of pink dashed rectangular regions shown in **b)**. NaN3 was added at 13 min (shown as orange line between 12.5 min and 17.5 min time points). Orange and pink arrows highlight response to NaN3 addition, resulting in inhibition of ATP synthesis through oxidative phosphorylation and triggering fragmentation and circularization of mitochondria. **d)** Representative maximum intensity projections of a 3-fold nematode embryo expressing GCaMP3 from the *nmr-1* promoter, highlighting calcium transient during backwards movement. Red circle: AVA cell body, green dashed line ventral nerve cord (VNC). **e-g** indicate higher magnification views of VNC, corresponding to green dashed rectangular regions in **d**. Red and yellow arrows indicate relative positions (posterior, anterior) along VNC. A kymograph of dF/F along the VNC is shown in **h**, diagonal white arrow emphasizes intensity gradient progressing along space and time. Intensity profiles corresponding to red rectangular region at top are further plotted in **i**, for reference, intensity derived from the AVA cell body is also shown. dF/F derived from the two views and their average are plotted. Grey box denotes period of backwards locomotion. **j)** Higher magnification view of yellow dashed rectangular region in **d.** Contrast has been adjusted (saturating cell bodies) in order emphasize fine spatial details within the embryonic nerve ring. All images are deconvolved, derived from direct and reflected 1.1 NA views. All perspectives are shown relative to the coverslip, except in **d**, where rotation angle relative to Y axis is shown. All scale bars: 5 μm, except in **c**, 1 μm. See also **Supplementary Videos 9-12.**

The polarization of the microtubule organizing center (MTOC) towards antigen presenting cells (APC) is a critical step in the formation of the immunological synapse (IS) by T cells^18-20^. During IS formation, the T cell nucleus undergoes significant deformations^21,22^. To visualize this process, we simulated IS formation by depositing Jurkat T cells transfected with 3XEGFP-EMTB^23^ on reflective coverslips coated with anti-CD3 antibody (mimicking the APC). After the T cells spread on the coverslip, we collected 100 imaging volumes (one volume every 4 s) spanning almost 7 minutes (**Supplementary Video 9**). Spatial resolution was sufficient to resolve microtubule bundles at the base of the cell and around the cell nucleus (Fig. 5a). Intriguingly, we observed correlated movement of the centrosome, associated microtubules (positioned near the coverslip), and inward deformation of the nucleus (4 μm away), suggesting that the movement of microtubule filaments and associated molecular motors around the centrosome may be involved in pushing and deforming the nucleus.

To visualize longer-term changes in response to a perturbation, we labeled mitochondria with GFP in U2OS cells, and added sodium azide 13 minutes after imaging commenced (Fig. 5b, **Supplementary Video 10**). Sodium azide inhibits cytochrome c oxidase, a key complex involved in oxidative phosphorylation. The resulting block on respiration caused a dramatic fragmentation and circularization of mitochondria, changes that were easily resolved with our system (Fig. 5c), over our 50- minute recording. As another example of our ability to visualize extended dynamics at the sub-organelle spatial scale, we visualized GFP-tagged galactosyltransferase, a resident protein of the trans-Golgi compartment, in U2OS cells, capturing volumes every 15 s for 45 minutes (**Supplementary Video 11**). High NA reflective imaging revealed highly dynamic, ribbon-like Golgi stacks juxtaposed around the nucleus, as well as rapidly moving Golgi vesicles.

In order to illustrate our system’s capability to image fine structures and rapid dynamics in a somewhat larger sample, we performed functional imaging in a freely moving, late-stage (3-fold) nematode embryo (Fig. 5d, **Supplementary Video 12**). For calcium imaging studies in *C. elegans*, fluorescent intensity is typically extracted from the cell body. This simplifies image analysis, particularly for freely moving animals, but it ignores signals compartmentalized in neurites^24^. Using a single-view iSPIM^15^ with 0.8 NA detection for functional imaging in *C. elegans* embryos, we recently demonstrated that calcium transients in the AVA cell bodies correlated with backward movement in the eggshell^16^. The improved collection efficiency, near-isotropic resolution, and high speed of the high-NA system allowed us to extend this work, as we were able to document calcium transients in neurites (Fig. 5d-g), as well as cell bodies. Using a strain expressing GCaMP3 from the *nmr-1* promoter^25^, we extracted fluorescence intensity from the cell body of reversal command interneuron AVA, forward command interneuron PVC, as well as the ventral nerve cord, which contained the neurites of both AVA and PVC (Fig. 5d). As reported previously^16^, backward movement in the eggshell positively correlated with activity in AVA (Fig. 5i), but not PVC (**Supplementary Fig. 7**), cell bodies. These calcium transients were also apparent in the ventral nerve cord, which showed an even more striking rise during backward motion (Fig. 5d). Our imaging rate of ^~^1.6 volumes/s was sufficiently fast to observe local dynamics along the length of the VNC (Fig. 5h). Since the transient was observed in the raw normal and mirrored views, it is highly unlikely that the intensity change was due to imaging- or post-processing artifacts. Finally, we note that the spatial resolution in our reconstructed images was sufficient to observe substructure within the embryonic nerve ring (Fig. 5j), detail that is normally masked by diffraction in conventional optical imaging.

## Discussion

Our method extracts additional information from the limited fluorescence photon budget, using reflection to improve the speed, resolution, and light collection efficiency of LSFM. We speculate that similar approaches could be applied fruitfully in super-resolution microscopy, which remains limited to single-objective collection. The commercially available reflective coverslips we used provided an economic method for adding extra views to the base LSFM system without introducing expensive objectives or significantly complicating instrument design due to the associated steric hindrance. The price that must be paid for these benefits is an increased computational burden, as more careful modeling of the imaging process and extensive post-processing are necessary to remove epifluorescence contamination within the raw data. In contrast to most previous efforts that apply shift-invariant deconvolution to LSFM, we developed a more accurate shift varying model of image formation, which is necessary to remove the spatially varying background in our data. The consequence is that for large datasets, processing time is still a practical bottleneck (**Supplementary Table 1**). Speeding up the processing, either using improved hardware (e.g. GPUs with more memory), parallelizing (e.g. more GPUs or many CPU cores), or by optimizing the code is an important next step in improving the technique.

The light sheet in the reflective diSPIM we present here is thicker than ideal, because the illumination in our reflective geometry must traverse twice as much sample as in conventional diSPIM that alternates illumination from each objective. We did not address this issue here, as we observed no significant deterioration in image quality in our modestly sized (thickness < 50 μm) samples. An interesting future direction might be to pair our method with approaches that engineer the excitation point spread function^26,27^ to create pseudo light sheets with effectively thinner width over the entire imaging field, thereby improving axial resolution. Such an approach could also be used to improve lateral resolution, if sharp structure is introduced within the image plane, although PSF engineering of any kind implies more dose (sidelobes) delivered outside the focal plane. Finally, we note that our approach could be adapted to use a single high NA objective for introducing direct and reflected light sheets and collecting the corresponding views, improving spatiotemporal resolution and signal collection and further simplifying instrument design (**Supplementary Note 3**).

## Acknowledgments

We thank Gary Rondeau and Jon Daniels (Applied Scientific Instrumentation) for help in designing the asymmetric diSPIM and useful discussion regarding motorized and piezoelectric stages, Henry Eden (NIBIB) and Talley Lambert (Harvard Medical School) for carefully reading the manuscript, Alan Hoofring (NIH Medical Arts Design Section) for help with illustrations, and Chris Janetopoulos (University of the Sciences) for interesting discussion regarding reflective imaging. This work was supported by the Intramural Research Program of the National Institute of Biomedical Imaging and Bioengineering at the National Institutes of Health. P.L. and H.S. acknowledge summer support from the Marine Biological Laboratory at Woods Hole, through the Whitman- and Fellows- program. P.L. acknowledges support from NIH National Institute of Biomedical Imaging and Bioengineering (NIBIB) of the National Institutes of Health (NIH) under grant number R01EB017293. C.S. acknowledges funding from the National Institute of General Medical Sciences of NIH under Award Number R25GM109439 (Project Title: University of Chicago Initiative for Maximizing Student Development [IMSD]) and NIBIB under grant number T32 EB002103. Partial funding for the computation in this work was provided by NIH Grant Nos. S10 RRO21039 and P30 CA14599. A.U. and I.R-S. were supported by the NSF grant number 1607645. The contents of this article are solely the responsibility of the authors and do not necessarily reflects the views of the NIH.

## Author Contributions

Conceived project: Y.W., J.T., H.S. Developed imaging model: C.S., Y.W., H.S., and P.L. Implemented imaging model: Y.W. and C.S., with guidance from P.L. Designed optical system: Y.W., A.K., and H.S. Built optical systems and wrote acquisition software: Y.W., A.K. and M.G. Acquired data: Y.W. Provided guidance on biological experiments and prepared samples: E.A., P.C., R.C., I.R-S., H.V., J.C., and A.U. All authors analyzed data. Wrote paper: Y.W., P.L, and H.S. with input from all authors. Supervised research: P.L. and H.S.

## Methods

### Sample preparation

#### Nematode embryos

*C. elegans* were maintained on nematode growth medium seeded with *Escherichia coli* (OP50). Embryos were dissected from gravid adults, placed on poly-L-lysine coated coverslips and imaged in M9 buffer, as previously described^15^. Strain BV24 [*ltIs44* [*pie-1*p-mCherry::PH(PLC1delta1) + *unc-119*(+)]; *zuIs178* [(*his*-72 1kb::HIS-72::GFP); *unc-119*(+)] V] was used for imaging nuclei (Fig. 1 and **Supplementary Videos 1-3**); strain AQ2953 *ljIs131*[*myo-3*p::GCaMP3-SL2-tagRFP-T]^28^ for imaging calcium flux within 3- fold embryos (Fig. 2 and **Supplementary Videos 4-5**); strain TQ3032 *xuEx1040*[*nmr-1p::GCaMP3; nmr-1p::DsRed*]^25^ for imaging calcium transients in neurons and neurites during backwards movement (Fig. 5d-i, and **Supplementary Video 13**); and strain SLS-5 [*jIS13* [*pie-1::mCherry::H2B + unc-119(+); pnhr2::mCHerry::histone + unc-119(+)*]*II, wIS51* [*pSCM::GFP + unc-119(+)*]*, mcIS50* [*lin-26p::vab-10 (actin binding domain)::GFP + myo-2p::GFP + pBluescript*]] was used for the comparison of conventional 0.8/0.8 NA diSPIM and 1.1/0.71 NA diSPIM (**Supplementary Fig. 5**).

#### Jurkat T cells

Reflective coverslips were rinsed with 70% ethanol, dried with filtered air, and incubated in 0.01% W/V Poly-L-Lysine (PLL, Sigma Aldrich, St. Louis, MO) for 10 minutes. PLL was aspirated and the coverslip left to dry for 1 hour at 37°C. Coverslips were coated with T cell activating antibody by incubating the coverslips in a 10 μg/mL solution of anti-CD3 antibody (Hit-3a, eBiosciences, San Diego, CA) for 2 hours at 37°C. Excess antibody was removed by washing with L-15 imaging medium immediately prior to imaging. E6-1 Jurkat T-cells were transiently transfected with EMTB-3XGFP plasmid (a kind gift from William Bement, Addgene plasmid # 26741) using the Neon (Thermofisher Scientific) electroporation system two days before the experiment. Cells were imaged within one hour after plating on the reflective coverslips (Fig.5a, **Supplementary Video 10)**.

#### U2OS cells

Human osteosarcoma U2OS cells were routinely passaged in DMEM (Gibco 10569-010) supplemented with 10% FBS (Hyclone) and penicillin/streptomycin at 37°C, with 5% CO2. The day prior to transfection, cells were plated on reflective coverslips, at a density of ~60%. Cells were transfected with the appropriate plasmid using Turbofect (Life Technologies) at a ratio of 3:1 (Liposomes:DNA). The next day, the medium was replaced with fresh DMEM plus 10% FBS without phenol red, which was also used as the imaging medium. To image the plasma membrane (Fig. 3) we used Lck-GFP (Origene, Cat. No. RC100017), for mitochondria (Fig. 3, Fig. 5b-c, **Supplementary Video 11**) we used either mApple-TOMM20-N-10 (Addgene, Cat. No. 54955) or pShooter pEF-Myc-mito-GFP (Clontech), and for the trans-Golgi compartment (**Supplementary Video 12**) we used EGFP-GalT (Addgene, Cat. No. 11929).

For imaging actin, cells were fixed with 4% Paraformaldehyde (PFA) in PBS for 10 min, washed with 1X PBS, treated with blocking buffer (1X PBS with Tween 20, 1% BSA and 0.1% Triton X-100) for 10 min, and stained with Alexa Fluor 488 Phalloidin (A12379, ThermoFisher) for 30 min before imaging.

For imaging microtubules, U2OS cells were grown on uncoated coverslips until ^~^50% confluency. The entire coverslip was then submerged for 3 minutes in methanol pre-chilled to -20°C to fix the cells. Coverslips were then washed in room temperature PBS extensively before blocking in antibody dilution buffer (Abdil; 1%BSA, 0.3% Triton-X 100 in PBS) for 1 hour at room temperature. The primary antibody stain was performed overnight at 4°C using 1/500 mg/ml of mouse anti alpha-Tubulin (Thermo Scientific #62204) in Abdil. The secondary antibody stain was performed for 1-2 hours at room temperature using 1/200 mg/ml of goat anti-mouse Alexa 488 (Invitrogen A11001) in Abdil.

#### Reflective coverslips

Reflective coverslips were purchased from Thin Film Coating, LLC (Breckenridge, MN). Coverslips were mirrored by sputtering a 150 nm thick aluminum film over the entire glass coverslip (24 mm × 50 mm × 0.17 mm for symmetric 0.8/0.8 NA diSPIM or 24 mm × 60 mm × 0.17 mm for asymmetric 1.1/0.7 NA diSPIM) and then protected with a 700-nm-thick layer of SiO2.

### Symmetric 0.8/0.8 NA diSPIM

The diSPIM frame, excitation optics and detection optics for reflective imaging are similar to those in our previously reported free-space coupled diSPIM^6^ except that an XY piezo stage (Physik Instrumente, P-545.2C7, 200 μm × 200 μm) was bolted on the top of a motorized XY stage (Applied Scientific Instrumentation, RAMM and MS-2000). Glass coverslips (24 mm × 50 mm, #1.5, VWR, 48393241) or reflective coverslips (24 mm × 50 mm, #1.5, Thin Film Coating LLC) containing samples were placed in an imaging chamber (Applied Scientific Instrumentation, I-3078-2450), then mounted to the piezo stage. The MS-2000 stage was used for coarse sample positioning before imaging, and the piezo stage was used to step the sample through the stationary light sheets to obtain imaging volumes.

Two 40x, 0.8 NA water immersion objectives (Nikon Cat. # MRD07420) were held in the conventional, perpendicular diSPIM configuration with a custom objective mount (Applied Scientific Instrumentation, RAO-DUAL-PI). Each objective was housed within a piezoelectric objective positioner (PZT, Physik Instrumente, PIFOC-P726), enabling independent axial control of each detection objective.

For excitation, a 488 nm laser (Newport, PC14584) and a 561 nm laser (Crystalaser, CL-561-050) were combined with a dichroic mirror (Semrock, Di01-R488-25x36), then passed through an acoustooptic tunable filter (AOTF, Quanta Tech, AOTFnC-400.650-TN) for power and shuttering control. Light sheets were created by digitally scanning a Gaussian beam with a galvo (Thorlabs, GVS001) and relaying the image of the galvo to the back focal plane of an excitation objective. Unlike conventional diSPIM imaging on glass coverslips where the light sheet beam waist is usually positioned at the center of the sample, the beam waist in reflective imaging was positioned at the surface of the coverslip.

For detection, direct fluorescence and mirror images were simultaneously collected via the two objectives, transmitted through dichroic mirrors (Chroma, ZT405/488/561/640rpc), filtered through long-pass and notch emission filters (Semrock, LP02-488RU-25 and NF03-561E-25) to reject 488-nm and 561-nm excitation light, respectively, and imaged with 200-mm tube lenses (Applied Scientific Instrumentation, C60-TUBE_B) onto two scientific-grade, complementary, metal-oxide-semiconductor (sCMOS) cameras (PCO, Edge 5.5). The image pixel size was 6.5 μm/40 = 162.5 nm, resulting in a field of view up to ^~^110 μm × 80 μm (e.g., in Fig. 1b-c).

In some experiments (Fig. 1, **Supplementary Fig. 2** and **Supplementary Video 4**), the cameras were operated in rolling-shutter mode, synchronized with the illumination line scan. This feature enabled partially confocal detection, improving background rejection^29^. The slit width was set at 1.6 μm to match the FWHM of the light-sheet beam waist. Note that in this mode, imaging is slower because the camera must read out unidirectionally, from the top row of the camera to the bottom row. To pursue higher-speed imaging in most experiments, we thus dispensed with the synchronized rolling shutter, instead using the default rolling mode (where detection is read out both top-to-bottom and bottom-to-top, thus improving imaging rate 2-fold over the synchronized rolling shutter mode).

### Asymmetric 1.1/0.71 NA diSPIM

The asymmetric diSPIM frame, excitation optics and detection optics are similar to those in our previous, fiber coupled diSPIM^17^. An XY piezo stage (Physik Instrumente, P-545.2C7, 200 μm × 200 μm) was bolted on top of a motorized XY stage (Applied Scientific Instrumentation, RAMM and MS-2500). Conventional glass coverslips (24 mm × 60 mm, #1.5, Electron Microscopy Sciences, Cat # 63793-01) or reflective coverslips (24 mm × 60 mm, #1.5, Thin Film Coating LLC) containing samples were placed in an imaging chamber (Applied Scientific Instrumentation, I-3078-2460), then mounted to the piezo stage. The MS-2500 stage was used for coarse sample positioning before imaging, and the piezo stage was used to step the sample through the stationary light sheets to create imaging volumes.

A 25x, 1.1 NA water immersion objective (Nikon, Cat. # MRD77220, f = 8 mm) and a 28.6x, 0.71 NA water immersion objective (Special Optics, f = 7 mm) were held in an objective mount (Applied Scientific Instrumentation, CDZ-1000). Because the 1.1 NA and 0.71 objectives have angular apertures ^~^111.6 and ^~^64.5 degrees, respectively, the two lenses cannot be assembled at 45 degrees relative to the vertical axis while still using a horizontally mounted coverslip, as in conventional diSPIM. Thus the objective mount was tilted ^~^12 degrees relative to the vertical axis so that the 1.1 NA and 0.71 NA objectives were rotated ^~^33 degrees and ^~^57 degrees relative to vertical axis, respectively (**Supplementary Fig. 4**). The objectives were housed within piezoelectric objective positioners (Applied Scientific Instrumentation, PZ, X-Y ADJUSTABLE), enabling independent axial control of each detection objective.

For excitation, 488 nm and 561 nm lasers (150 mW each, Coherent, OBIS model, 1277611 and 1280720) were combined with a dichroic mirror (Semrock, Di01-R488-25x36), then passed through an acousto-optic tunable filter (AOTF, Quanta Tech, AOTFnC-400.650-TN) for power and shuttering control. Light sheets were created by digitally scanning a Gaussian beam with a MEMS mirror (Applied Scientific Instrumentation, Tiger controller, TG1000) that is conjugate to the back focal plane of the objectives. The beam waist was set at 1.6 μm thickness (FWHM value) and focused at the coverslip (i.e., the center of the sample and its mirror image).

For detection, direct fluorescence and mirror images were collected via the two objectives, transmitted through dichroic mirrors (Chroma, ZT405/488/561/640rpc) and a multiple band-pass filter (Chroma, 59022m), and imaged with 400 mm tube lens (for the 1.1 NA detection path, Applied Scientific Instrumentation, C60-TUBE_400) and 200 mm tube lens (for the 0.71 NA detection path, Applied Scientific Instrumentation, C60-TUBE_B) onto sCMOS cameras (Hamamatsu, ORCA Flash 4.0 v2), respectively. The effective magnifications for 1.1 NA and 0.7 NA detection were 50x and 28.6x, respectively; so that the resulting image pixel size was 6.5 μm/50 = 130 nm for the high NA view, and 6.5 μm/28.6 = 227 nm for the low NA view, respectively (resulting in a field of view up to ^~^130 μm × 130 μm, e.g. in Fig. 4g).

### Data Acquisition

The control waveforms for implementing single- and dual-color 0.8/0.8 NA reflective diSPIM imaging are similar to those in our previous triple-view light sheet imaging^1^, except that excitation was usually introduced from one objective (i.e., we did not use alternating excitation, except for the comparative 0.8 NA/0.8 NA dual-view images and 1.1 NA/0.71 NA quadruple-view images shown in Fig. 4d). The waveforms include a step-wise waveform (e.g., 200 Hz/plane, 250 ms volume imaging rate for 50 planes) to drive the XY piezo stage, and two identical external trigger signals to simultaneously trigger the two sCMOS cameras. Programs controlling DAQ waveforms were written in Labview (National Instruments) and programs controlling image acquisition (via PCO sCMOS cameras) were written in the Python programming language.

The control waveforms for implementing single-color 1.1/0.71 NA reflective diSPIM imaging are similar to those used in our fiber-coupled diSPIM^14^, and include a step-wise waveform to drive the XY piezo stage and two external trigger signals to sequentially trigger the two sCMOS cameras. Programs controlling DAQ waveforms and Hamamatsu sCMOS image acquisition were both written in Labview (National Instruments). All software programs are available upon request. **Supplementary Table 2** summarizes data acquisition parameters used in this work.

### Data Processing

Raw image data from the four views in 0.8/0.8 NA reflective imaging or two views in 1.1/1.1 NA reflective imaging are merged to produce a single volumetric view, after processing steps that include background subtraction, interpolation, transformation, fusion, registration, epifluorescence removal and joint deconvolution. Background subtraction and interpolation procedures are described elsewhere^1^. When processing symmetric diSPIM reflective data, the transformation converts the raw data acquired in stage-scanning mode to light-sheet scanning mode, i.e. viewed from the objective, as if the stage were stationary and the light sheet moved through the sample^6^. In the asymmetric configuration, raw data were further transformed so that they are viewed from the perspective of the coverslip^1^.

Registration is implemented in the open-source Medical Imaging Processing, Analyzing and Visualization (MIPAV) programming environment (http://mipav.cit.nih.gov/), other processing steps are implemented in MATLAB (R2015a) with both CPU (Intel Xeon, ES-2690-v3, 48 threads, 128GB memory) and GPU (using an Nvidia Tesla K80 graphics card, 2 GPU units, each 12GB memory) programming. After transformation to light-sheet scanning perspective, the volumetric views are combined to one view, and then virtually mirrored to form a second view (see the subsection *‘Fusing raw volumetric views to create two views’* and Equation 10 for this procedure). The two views are then registered^1^ and jointly deconvolved with a new method that simultaneously removes spatially varying epifluorescence contamination and recovers the best spatial resolution inherent to each raw view. Below we describe the theoretical concept and implementation of this joint deconvolution algorithm by (i) modeling the image formation process in reflective imaging; (ii) deriving the deconvolution method for the removal of spatially varying epifluorescence contamination in single-view imaging; (iii) fusing raw volumetric views to create two complementary views, each with anisotropic spatial resolution; and (iv) implementing joint deconvolution of the two views to remove epifluorescence contamination as well as resolution recovery.

### *1.* Image formation process in reflective imaging

We start by modeling image formation in light-sheet scanning mode, i.e. moving the light sheet and its reflection through the stationary samples. Using a fixed coordinate system *x,y,z* (**Supplementary Fig. 8)** and assuming the mirror lies in the plane *z* = −*x* tan(θ) (e.g., *z* = −*x* for the 0.8/0.8 NA symmetrical configuration as θ = 45°), we can model the effect of the mirrored coverslip by reflecting the object fluorophore distribution *f*(*x, y, z*) across the coverslip to obtain an object distribution:

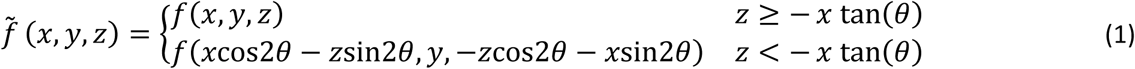

The illumination pattern is denoted *I* (*x, y, z*) and is the sum of the incident excitation sheet *I*_1_, its direct reflection *I*_2_ and their mirror images *I*_3_ and *I*_4_:

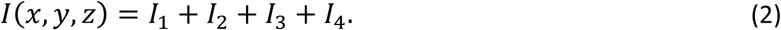

We assume that the illumination pattern moves along the mirror by an increment *δ* in the *x* direction and an increment −*ξ* in the *z* direction, where *ξ* **=** *δ* tan(θ). This illumination shift relative to the fixed coordinate system can be represented as:

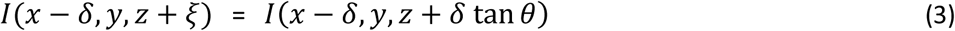

Consider a camera looking down the *z* axis, recording an image in camera coordinates *x*′, *y*′. If we assume the objective has a response function *h*(*x, y, z*) that is shift invariant in *x* and *y* and focused at the plane *z* **=** −*δ* tan *θ*, then the image recorded by the camera at a given illumination position *δ* can be written

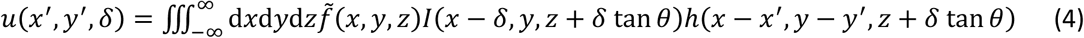

By comparing this image-formation formula for the reflective imaging case with that derived for imaging on conventional coverslips (**Supplementary Note 1),** we find that Equation 4 can be decomposed into two terms: one represents conventional in-focus imaging of the light sheet (assumed to be shift invariant), and the other term represents a spatially variant epifluorescence contamination.

To further model the image-formation and deconvolution process, we consider a discrete formulation, in which the acquired three-dimensional image stack is reformulated as a one-dimensional vector ***u*** with a total of voxels *u_i_, j* = 0,…, J − 1. The goal is to determine from these measurements the object fluorophore distribution 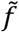, also represented as a one-dimensional vector 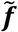 of K voxels 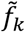, *k* = 0,…, K − 1. Because the imaging equation (4) is linear, the mapping between the vectors 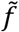 and *u* can be expressed as

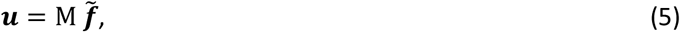

where M is a *J* × *K* matrix. As we will discuss below, we do not explicitly construct the matrix M*_δ_*, but it has a block structure comprising blocks of rows, which we denote M*_δ_*, corresponding to each illumination offset. Each such block could be constructed through multiplication of three matrices, M*_δ_* **=** PHD*_δ_*, modeling three cascaded operations: (1) a diagonal matrix D*_δ_* representing multiplication of the object fluorophore vector by the *δ*-shifted crossed light sheets; (2) a matrix H that is block circulant with circulant blocks that represents looping over *z* in the object distribution and performing 2D convolution with a slice of the detection objective point spread function *h*(*x, y, z*) at each *z*, and (3) a projection matrix P that models collapsing over z. Because of the spatially varying illumination, the matrix M is not shift invariant (i.e., it does not have a circulant or block-circulant structure).

### *2.* Single-view deconvolution to remove epifluorescence contamination

To recover the object distribution 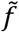 (i.e., including the true object *f* and its mirror image), the computational task is to invert Equation 5 (i.e., recovering object vector 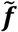 from the measured data vector ***u***). We achieve this using an iterative Richardson-Lucy update, which is an appropriate deconvolution method for our data, as the data are corrupted by Poisson noise.

For a general matrix *M* that is not necessarily shift-invariant or normalized to 1, the single-view Richardson-Lucy iterative update for the current estimate 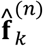 of the object distribution 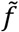 at voxel *k* and iteration *n* is given by^30^

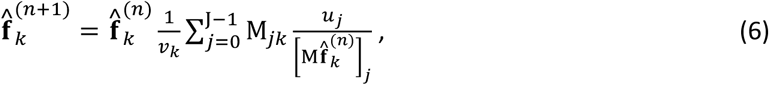

where 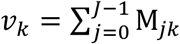 is the *k*th element of a sensitivity or normalization vector that ensures every iterative update is normalized. If we define a ratio vector **r** with elements 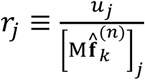, then the sum in Equation 6 is equivalent to multiplying the vector **r** by **M*^T^*** (where ***T*** denotes transpose):

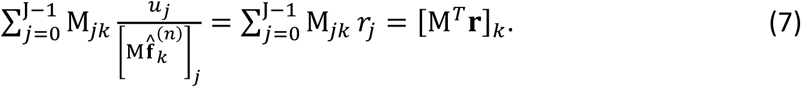

Note that the matrix transpose M^T^ maps from data space to the object space. Substituting Equation 7 into Equation 6, we can write the single-view Richardson-Lucy iterative update compactly as:

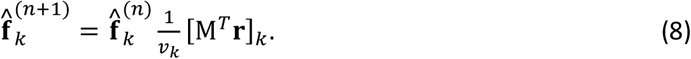

Note that M^T^ also has a block structure, with column blocks given by **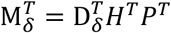**. Here P^T^ is a backprojection matrix, which smears the vector to which it is applied back across the image grid. *H^T^* represents looping over *z* in the object distribution and performing 2D convolution with a slice of the transposed detection objective point spread function at each *z*. Finally 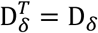 since it is diagonal; its action denotes multiplication with the shifted illumination pattern.

In practice, we do not explicitly construct the matrix *M* or convert the image stacks to one-dimensional vectors. We implement the Richardson-Lucy update by acting directly on the image stacks according to the operations described above. By changing the one-dimensional vectors back to image stacks (e.g. replace vectors 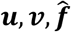 with image stacks that we denote 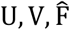, respectively), we can rewrite the above Richardson-Lucy iterative update (Equation 8) as:

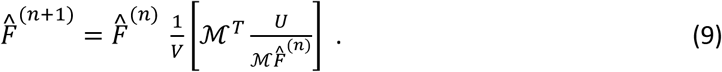

We now use script 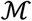 to denote the operator acting on these image stacks since it is not strictly a matrix. Equation 9 implies 4 steps in each RL-update: (1) Compute 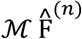 by applying the operator 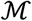 to the current estimate of the object 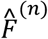 according to the three cascaded operations 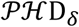 at each *δ* as described above; (2) divide the measured data stack *U* by this quantity, and denote the resulting ratio image 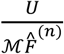 as *R*; (3) apply the transpose operator 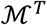 to *R*, which involves applying the cascaded operations 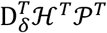 described above and then summing over the various shifts *δ*; (4) update the current estimate 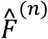 by multiplying by the correction image 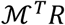 and dividing by the normalization image *V*. Note that from this point of view, the normalization image 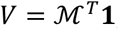, where **1** denotes an image of ones. More detailed discussion can be found in the **Supplementary Note 1.**

Applying the generalized single-view Richardson-Lucy deconvolution described above (Equation 9), the object distribution 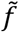 can be recovered, and thus epifluorescence contamination can be removed. The next step is to adapt this method for multiview imaging.

### *3.* Fusing raw volumetric views to create two views

In order to combine the benefits of epifluorescence removal while obtaining a more isotropic PSF, we form view *U***_1_** and a second, virtual view *U***_2_** by reflecting view U_1_ along the mirror (**Supplementary Fig. 9**). *U*_1_ and *U*_2_ are defined as follows:

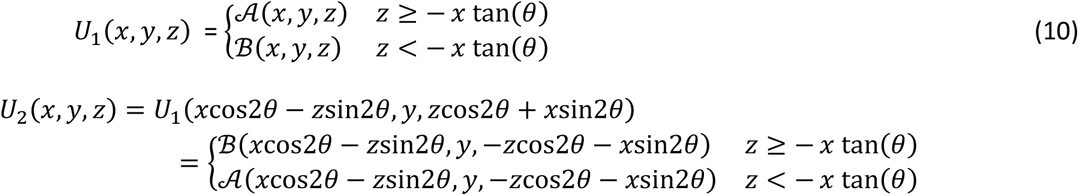

Here *U*_1_ is decomposed into two parts, 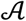 and 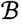, which are defined according to the imaging geometry. In the symmetric 0.8/0.8 NA diSPIM case where we simultaneously acquire 4 views (A, A’, B, and B in Fig. 1), *U*_1_ can be formed as the simple sum of views, i.e., 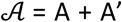 and 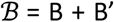 if all the 4 views have similar image quality. Alternatively, *U*_1_ is defined as the combination of data with best image quality, i.e., 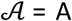 or A’ and 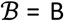 or B’ (**Supplementary Fig. 1)**. Note that before combining raw views, they must be registered (e.g. A with A’, B with B’). If there are only two views acquired from one camera (e.g., dual 1.1 NA view acquisition in the reflective 1.1/0.71 NA diSPIM system), *U*_1_ is the acquired image (A + B’), i.e., 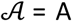; 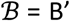.

Although 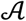 and 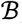 already contain complementary spatial frequencies, we treat them as a single image *U*_1_ visualized from a single perspective so that the epifluorescence contamination can be removed with the deconvolution procedure developed in the previous section. The virtual, reflected image of *U*_1_, i.e. *U*_2_, is visualized from a direction complimentary to *U*_1_. *U*_1_ and *U*_2_ are thus blurred with complementary detection PSFs (**Supplementary Fig. 9**), so they can be jointly deconvolved to simultaneously remove epifluorescence contamination and recover spatial resolution.

### *4.* Joint deconvolution for epifluorescence removal and resolution recovery

We register the two views *U*_1_ and *U*_2_, and perform joint deconvolution on them by applying the generalized Richardson-Lucy update described above to each view as follows:

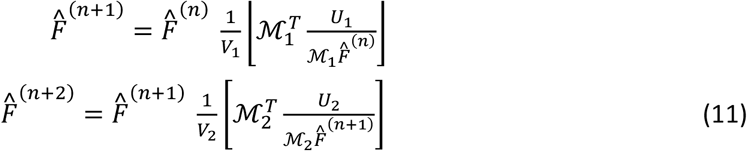

Here, 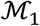 and 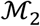 are the operators that map the object stack 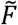 to either measured view stack *U*_1_ or virtually reflected view stack *U*_2_, respectively, and 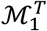 and 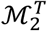 are the transpose operators that map from data space to object space. These are all implemented as in the single-view case described above, with the appropriate detection PSF for each view applied in the convolution operations 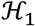 and 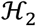 and the projection and backprojections applied over the view-appropriate axis. In the absence of rolling-shutter mode, the PSFs are simply the wide-field PSFs of the detection objectives, which are derived from the ‘Born and Wolf’ model with appropriate numerical aperture (**Supplementary Note 2**). When applying the rolling-shutter mode, we model the effect of the shutter by using an effective PSF that is the product of the wide-field PSFs with a slit function (**Supplementary Note 2**). *V*_1_ and *V*_2_ are the normalization images for the two views, respectively. 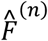 is the current estimate, 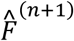 is the estimate derived from *U*_1_, and 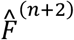 is the new estimate derived after considering *U*_2_.

Recall that the estimate of the object distribution 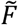 is a combination of the object *f*(*x, y, z*) and its reflection. Since *U*_1_ and *U*_2_ are also mirror images, joint deconvolution ensures that the estimate of either the object or its reflection are identical as convergence is reached. Therefore, we are free to pick either as our final estimate of the object *f* (*x*, *y*, *z*).

For 1.1 NA/0.7 NA quadruple-view imaging on reflective coverslips (Fig. 4d), *U*_1_ is defined as the acquired image (A + B’) with the 1.1 NA objective, i.e., 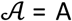; 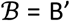. We create *U*_2_ as the reflection of *U*_1_, defined according to Equation 10. We define the image acquired with 0.71 NA objective as *U*_3_ and its reflected image as *U*_4_. These 4 image sets are alternately deconvolved by extending Equations 11 to a quadruple-view case, the same procedure we used for previous triple-view deconvolution^1^.

When applying naïve deconvolution for diSPIM imaging on reflective coverslips (as in Fig. 1e), we performed conventional joint deconvolution^6^ (using only the detection PSF of each lens) on the two quantities 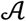 and 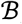 in *U*_1_. For diSPIM imaging on conventional glass coverslips (e.g., Fig. 4c, f, **Supplementary Fig. 6**), alternating deconvolution was performed on the two measured views as previously described^6^.

The above calculations (Equations 11) were implemented in MATLAB (version 2015a, with imaging processing toolbox and parallel computation toolbox) by employing a graphics processing unit (GPU, Nvidia, Tesla K80) to speed deconvolution of the registered volumes (*U*_1_, *U*_2_) (**Supplementary Software** and **Supplementary Note 1**). Note that in the calculation of forward operator 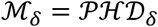 performing 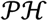 (i.e., 2D convolution at each z plane with the corresponding detection objective point spread function, and then collapsing over all z) is equivalent to 3D convolution at the axial plane z = 0; in the calculation of backward matrix 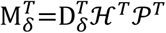, performing 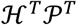 (i.e., back projection, and 2D convolution with the corresponding PSF at each z plane) is equivalent to 3D convolution of the data at z = 0 (i.e., creating a 3D stack with the values at z = 0 and zeros at other z) with the 3D PSF. By performing these 3D convolutions, processing time is greatly reduced as only 4 (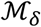 and 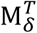 for each view) 3D convolutions (compared to 1440 2D convolutions for a stack with 360 z planes) at each stage scanning step (i.e., at each increment of *δ*) are required for each Richardson-Lucy update. However, the overall computation is still time consuming. For example, 20,000 3D convolutions are required for an imaging volume with 100 steps and 50 iterations of our modified deconvolution procedure; by contrast, only 200 3D convolutions are required in conventional joint deconvolution for the same number of iterations^6^. As before^1^, we used the NVIDIA *CUDA* fast Fourier transform library (*cuFFT*) with a GPU card (Tesla K80) to implement 3D convolutions in the Fourier domain, speeding processing. For example, for a stack size of 360 × 360 × 100 voxels, GPU processing required 90 s for one iteration, but a dual-CPU architecture (2.6 GHz, 24 cores, 48 threads, and 128GB memory) takes ^~^20 minutes for one iteration. Processing times for all datasets are listed in **Supplementary Table 1**, the number of iterations used for deconvolving each dataset are listed in **Supplementary Table 2**. As before^1^, we determined the number of iterations for a given dataset by empirically inspecting convergence based on image quality. In practice this number varied, depending on SNR (higher SNR allows more iterations before noise is amplified).

### Estimating spatial resolution

Optical transfer functions (OTFs) and spatial resolution estimates presented in **Supplementary Table 1** and Fig. 4c are based on simulations. For conventional LSFM imaging on glass coverslips, we simulated the PSF by blurring a single point with the detection point spread functions appropriate for the imaging geometry (0.8/0.8 NA or 1.1/0.71 NA) with an ImageJ plugin (http://bigwww.epfl.ch/algorithms/psfgenerator/), then multiplied it with the excitation light sheet (assuming a FWHM of 1.6 μm). For reflective imaging, we simulated a bead 1 μm away from the reflective coverslip, mirrored it, and then blurred it according to Equation 4. Joint deconvolution was performed on the simulated bead as previously reported for conventional imaging^1^, or with Equations 9-10 above for reflective imaging. In all cases, we used 10 iterations to achieve convergence for x’ and z’ dimensions. However, we noticed that convergence in the y’ dimension was faster than either x’ or z’. For example, when the FHWM value in x’ was ^~^260 nm after 10 iterations (in the 1.1 NA case), the FWHM value in the y’ dimension dropped to ^~^215 nm, presumably beyond the diffraction limit. We suspect the over-optimistic convergence value in y’ arises due to intrinsic asymmetries in the FHWM values (e.g., 305 nm × 305 nm × 909 nm for 1.1 NA view; 472 nm × 472 nm × 2200 nm for 0.71 NA view), in which the difference of the initial FHWM values along y’ (i.e., 305 nm vs. 472 nm) is much less than the difference between x’ and z’ (i.e., 305 nm vs. 2200 nm). To circumvent this issue, we implemented post-smoothing (a well-known procedure in nuclear medicine image reconstruction^31,32^) along y’ so that the final FHWM value in y’ is identical to the FWHM in x’.

The kx/kz OTF cross sections derived from experiment (**Supplementary Fig. 5**) were produced by computing the 3D Fourier transform of the microtubule samples (Fig. 4d), then averaging all kx/kz cross sections along the ky dimension.

### Calculation of collection efficiency

The collection efficiencies in different objective/reflective configurations in **Supplementary Table 1** were calculated by computing the ratio of summed solid angles from all detection objectives to the total solid angle (4*π* steradians). The solid angle of each objective lens was calculated as 2*π*(1 − cos(*θ*)), where *θ* is half of the angular aperture, i.e., arcsin (NA/*n*), NA is the numerical aperture of the lens, and *n* is the refractive index of water.

### Image intensity correction

For some datasets (mitochondria in Fig. 3, **Supplementary Video 5**; microtubules in Fig. 5a, **Supplementary Video 9**), we performed standard bleaching correction based on an ImageJ Plugin (https://imagej.net/Bleach_Correction).

In some experiments, we observed a slight line artifact, obvious in maximum intensity projections of the raw data (Fig. 3h and **Supplementary Video 6**), which we attribute to residual epifluorescence contamination. With coarse sampling along the direction of stage scanning (e.g., the stage is stepped at every 1 μm in Fig. 3), the epifluorescence is poorly sampled (appearing discontinuous in the x’ direction, when visualized from the view of objective), and is almost--but not completely--removed by our algorithm.

In other experiments (Fig. 4g), we observed slight striping along the y’ direction, due to inhomogeneous absorption, which can in principle be lessened with an antistriping scanner^33^. Here we simply minimized the visual appearance of the artifact by dividing the raw data with the average of intensity along the y’ axis.

### Code availability

Deconvolution scripts in MATLAB are provided as **Supplementary Software.** LabVIEW control software is available from the corresponding author upon request.

### Data availability

The data that support the findings of this study are available from the corresponding author upon reasonable request.

